# Column subset selection for single-cell RNA-Seq clustering

**DOI:** 10.1101/159079

**Authors:** Shannon R. McCurdy, Vasilis Ntranos, Lior Pachter

## Abstract

The first step in the analysis of single-cell RNA sequencing (scRNA-Seq) is dimensionality reduction, which reduces noise and simplifies data visualization. However, techniques such as principal components analysis (PCA) fail to preserve non-negativity and sparsity structures present in the original matrices, and the coordinates of projected cells are not easily interpretable. Commonly used thresholding methods avoid those pitfalls, but ignore collinearity and covariance in the original matrix. We show that a deterministic column subset selection (DCSS) method possesses many of the favorable properties of PCA and common thresholding methods, while avoiding pitfalls from both. We derive new spectral bounds for DCSS. We apply DCSS to two measures of gene expression from two scRNA-Seq experiments with different clustering workflows, and compare to three thresholding methods. In each case study, the clusters based on the small subset of the complete gene expression profile selected by DCSS are similar to clusters produced from the full set. The resulting clusters are informative for cell type.

## 1. INTRODUCTION

Advances in RNA sequencing technology have recently made it possible to measure the genomewide expression profile of single cells (Tang *et al*., 2009). This promising technology is not without computational and analytical challenges, some of which include quality control, quantification, normalization, technical variability, and other confounding factors such as batch effects (Stegle, Teichmann and Marioni, 2015; Wagner, Regev and Yosef, 2016). More general challenges stem from the high dimensionality of the expression profiles: for example, selecting informative features from within the expression profiles.

One use for single-cell RNA sequencing (scRNA-Seq) data is the characterization of heterogeneity of expression within a population of cells and the discovery of new cell types through clustering of expression profiles (Zeisel *et al*., 2015). This note explores the following question: is it possible reduce the number of features in the expression profile without a large effect on the error rate for clustering and classification? This question is inspired by the quality control and technical variability challenges of scRNA-Seq. Common techniques for quality control and technical variability reduction include simple thresholding schemes and principal components analysis (PCA). Both of these techniques reduce the number of features in the data matrix.

One commonly used technique to reduce the number of features in the data matrix involves selecting columns from the original data matrix **A**, to form a column submatrix **C**, by thresholding the individual columns based on a score. Frequently used scores are on measures of abundance (Lun, McCarthy and Marioni, 2016), empirical variance (Kwon, Fan and Kharchenko, 2017), abundance and empirical variance (McCarthy *et al*.), and index of dispersion (empirical variance/mean) (Satija *et al*., 2015; Trapnell *et al*., 2014). Read count thresholds are intended to reduce low-abundance genes (Bourgon, Gentleman and Huber, 2010) or genes with high dropout rates (Brennecke *et al*.,2013), as these genes are not considered informative. Variance thresholding methods assume that the most variable genes are responsible for the important differences between cells (McCarthy *et al*.). Index of dispersion thresholding has a natural interpretation in terms of formal hypothesis testing, when the null model for gene abundance is the Poisson distribution (Cox and Lewis, 1966). We call these methods *simple* thresholding methods, because the score for each column *i* depends only on column *i*. Furthermore, within each column *i*, covariance between the rows (cells) of that column is not taken into account. By selecting columns and not linear combinations of columns from **A**, the elements of **C** will maintain the properties of non-negativity, sparsity, and interpretability, an advantage over PCA, but there are no guarantees that **C** will have similar properties to the original data matrix **A**.

Replacing the original data matrix of scRNA-Seq expression profiles with a rank-*k* PCA truncation of the profiles is another commonly used technique to reduce the number of features and the technical variability (Wagner, Regev and Yosef, 2016). To understand the PCA truncation, we must establish some matrix notation that we will use throughout this note. We orient the original data matrix **A** so that the n rows are cells and d columns are features, where *n < d.* For PCA, singular value decomposition (SVD) is performed on the column-mean centered matrix **Ã = A − 1***μ*^*T*^, where **1** is an *n* × 1 column vector and 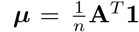 is a *d* × 1 column vector of column-means. The sum of the spectrum of eigenvalues of **ÃÃ**^*T*^ is proportional to the total empirical variance of **A**. The rank-*k* PCA truncation of **A**, which we call 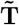, is the rank-*k* SVD truncation of **Ã**. SVD is reviewed in Sec. 6.1, and the formula for 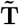 is provided there. As a consequence of the SVD, the spectrum of the square of the rank-*k* PCA truncation 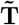 is identical to the spectrum of the square of the mean-centered data matrix **Ã** up to rank *k*; PCA gives a rank-*k* approximation to the mean-centered data **Ã** that preserves the maximum empirical variance of **A**. PCA is performed to reduce technical variability under the assumption that the technical variation is primarily captured by the non-leading eigenvalues and eigenvectors of **ÃÃ**^*T*^.

The drawback of replacing the original data matrix with the rank-*k* PCA truncation of the data that it fails to preserve non-negativity and sparsity structures present in the original data matrix, and the coordinates of projected cells are not interpretable in terms of single features.

The goal of column subset selection (CSS) is to extract from a matrix **A** a column submatrix **C** that conserves favorable properties, such as conditions on the spectrum of the column submatrix **C** (Tropp, 2009). Like the simple thresholding methods, CSS maintains the properties of nonnegativity, sparsity, and interpretability, and like PCA, CSS conserves favorable matrix properties. Similar to the simple thresholding methods discussed above, each column has a score, however in CSS algorithms, the score for each column *i* also depends on all of the other columns. We will consider rank-*k* subspace leverage scores in this note. Leverage scores have been considered for regression diagnostics and outlier detection in statistics (Velleman and Welsch, 1981; Chatterjee and Hadi, 1986) and were brought to prominence more recently in the context of randomized matrix algorithms (Drineas, Mahoney and Muthukrishnan, 2006). The rank-*k* subspace leverage score 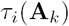 for the *i^th^* column of **A** is,

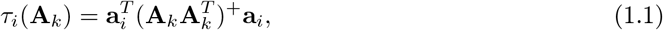

where the *i^th^* column of **A** is an (n × 1)-vector denoted by a_*i*_, **M**^+^ denotes Moore-Penrose pseudoinverse of **M**, and **A**_*k*_ is the rank-*k* SVD approximation to **A**, defined in Sec. 6.1. The leverage score 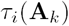 can also be written as the solution to the following optimization problem (Cohen *et al*., 2015),

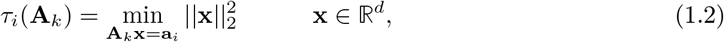

where 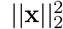 refers to the Euclidean (*L*_2_) norm of the vector x. The vector x measures how easily the column *a_i_* can be written as a linear combination of the columns of **A**_*k*_. Eqn. 1.2 shows that leverage scores capture the importance of each column a_*i*_ in the column space of **A**_*k*_ and are sensitive to collinearity between columns. We illustrate this point with a toy example in Sec. 2.1.

CSS algorithms select columns either with a random sampling procedure (such as in Drineas, Mahoney and Muthukrishnan (2006)) or a deterministic procedure. We showcase the deterministic CSS (DCSS) algorithm introduced by Papailiopoulos, Kyrillidis and Boutsidis (2014). Papailiopoulos, Kyrillidis and Boutsidis (2014) show that for datasets with power-law decay in 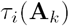, DCSS will select a least-squares approximation for **A**, **CC**^†^**A**, requiring fewer columns with the same accuracy than random sampling methods. One of the contributions of this note is a new bound for the spectrum of the square of **C** selected by DCSS projected onto the rank-*k* subspace that best approximates **A** (Eqn. 2.9). This bound means that, once both **C** and **A** are projected onto the rank-*k* subspace that best approximates **A**, **CC**^*T*^ is “close" to **AA**^*T*^. Another consequence is that the Frobenius norm of **C** is bounded (Eqn. 2.10). The Frobenius norm is a measure of the “size" of a matrix, so this bound provides confidence that the DCSS column matrix **C** is also similar in “size" to **A** and **A**_*k*_. In the event that DCSS is performed on a mean-centered matrix **Ã**, the Frobenius norm provides a measure of empirical variance. We also show a similar bound holds for random sampling (Eqn. 2.11), and under the assumption of power-law decay, DCSS requires fewer columns for the same error than random sampling.

In addition to the spectral bound, we present two case studies on two different scRNA-Seq experimental and analysis workflows to illustrate empirically the effect of thresholding features with DCSS compared to read count, variance, and index of dispersion on clustering and classification. To the best of our knowledge, this is the first time DCSS has been applied to scRNA-Seq data. The first case study is the genome-wide expression profiles of 3, 005 cells from the mouse cortex and hippocampus (Zeisel *et al.,* 2015) and the clustering workflow of Ntranos *et al.* (2016). The second case is the genome-wide expression profiles of 4, 423 cells from mouse bone marrow (Paul *et al.,* 2015) and the trajectory workflow of Setty *et al.* (2016). In both case studies, DCSS reduces the low abundance genes and maintains many of the most variable and over-dispersed genes. This shows that DCSS shares the best features of the simple thresholding methods and, like PCA, comes with additional bounds on the spectrum. This supports our conclusion that DCSS can be used instead of the simple thresholding methods for quality control and to reduce technical variability, in addition to selecting informative features. In both case studies, only a small fraction of the features are necessary to obtain clusters reflecting cell types, consistent with results in (Kwon, Fan and Kharchenko, 2017). We show that the error rate between the clustering assignments computed with the complete expression profile and the reduced expression profile is small.

## 2. METHODS

The aim of this note is to explore the effect of thresholding features, measurements of gene expression, with DCSS. We compare DCSS to simple thresholding methods and also to the complete data. These thresholding methods are the first step in the pre-processing workflow. In this section, we include the DCSS algorithm for completeness, and we describe the new bounds for DCSS.

### 2.1 The DCSS algorithm (Papailiopoulos, Kyrillidis and Boutsidis, 2014)

#### Algorithm 1.

*The DCSS algorithm selects for the submatrix **C** all columns i with a rank-k subspace leverage score 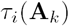 above a threshold θ, determined by the error tolerance ϵ and the rank, k. The algorithm is as follows.*

1. *Choose the rank, k, and the error tolerance, ϵ.*
2. *For every column i, calculate the rank-k subspace leverage scores 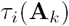 (Eqn. 1.1).*
3. *Sort the columns by 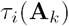, from largest to smallest. The sorted column indices are π_i_.*
4. *Define an empty set Θ = {}. Starting with the largest sorted column index π_0_, add the corresponding column index i to the set Θ, in decreasing order, until,*

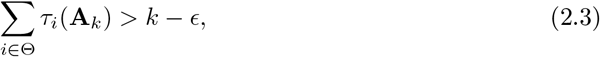

*and then stop. Note that 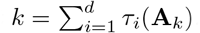. It will be useful to define 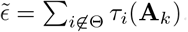. Eqn. 2.3 can equivalently be written as *ϵ* > 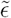.*
5. *If the set size |Θ| < k, continue adding columns in decreasing order until |Θ| = k*.
6. *The leverage score 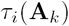 of the last column i included in Θ defines the leverage score threshold θ*.
7. *Introduce a rectangular selection matrix **S** of size d × |Θ|. If the column indexed by (i, π_i_) is in Θ, then 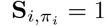. 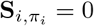 otherwise. The DCSS submatrix is **C** = **AS**.*

Theorem 3 of Papailiopoulos, Kyrillidis and Boutsidis (2014) states that when the rank-*k* subspace leverage scores exhibit a power-law decay in the sorted column index *π_i_*,

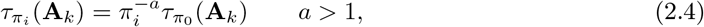

the number of sample columns selected by DCSS is,

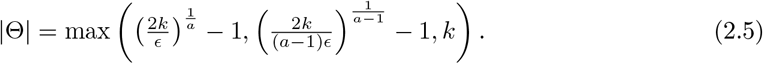

Papailiopoulos, Kyrillidis and Boutsidis (2014) demonstrate the power-law decay behavior of many real-world datasets; we show that this behavior is a reasonable assumption for the scRNA-Seq applications in Sec. 3.

For a statistical interpretation of DCSS, consider the data a_*i*_, *i* = 1,…, *d* to be identically and independently distributed (i.i.d.) according to the degenerate multivariate distribution 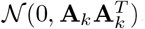. See Rao (1973) pg. 527-528 for a discussion of the degenerate multivariate distribution. Then the total likelihood of the data matrix **A** is,

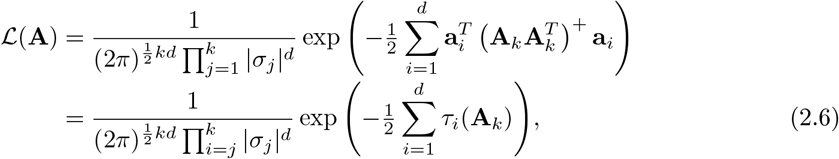

where |*σ_j_*| are the *k* largest singular values of **A**_*k*_. In contrast, the total likelihood of the DCSS matrix **C** is,

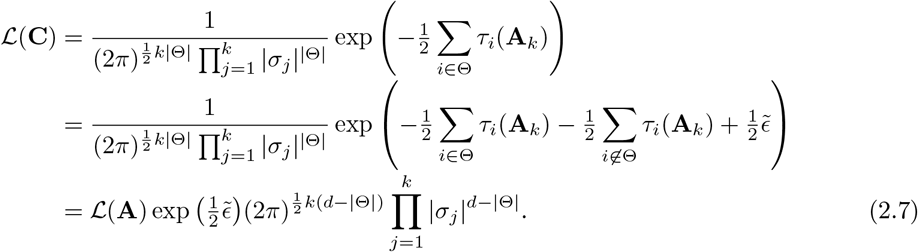

This shows that the DCSS matrix **C** preserves the total likelihood of the data up to a factor of 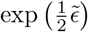 < 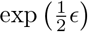 and a normalization constant, under the assumption that the data is i.i.d. according to 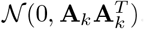. Any other selection set Θ′ of the same number of columns (|Θ′|=|Θ|) will have equal or greater error (*ϵ* ≤ *ϵ′*). This interpretation illustrates that DCSS accounts for covariance 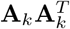between rows (cells). In contrast, the Poisson null model for the index of dispersion assumes independence between rows (cells) for each column (gene).

The DCSS method has two parameters, *k*, *ϵ* which jointly determine the number of columns |Θ| in the DCSS column submatrix **C**. The parameter *k* determines the rank of interest of the SVD approximation to **A**. The tuning parameter *ϵ* is a measure of the error tolerance in the “size" of **C** compared to **A_*k*_**. The selection of these parameters is a model selection problem, and in concert with a loss function, one could select these parameters using one’s preferred model selection method (e.g. cross-validation). The aim of this note, to compare clustering performed with the complete data matrix and a column submatrix, does not have a well-defined loss function, and so we use the heuristic “elbow" method for selecting *k* (Jolliffe, 2002), and we choose *ϵ* to be 0.1 or 0.05 in our applications.

As a toy example to illustrate how DCSS differs from the simple thresholding methods, consider the following toy data matrix,

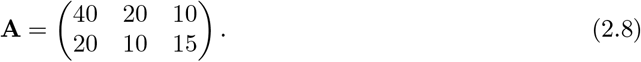

If the goal is to select a column submatrix with two columns, it is easy to check that simple thresholding by mean, variance, and index of dispersion all select the first and second columns. However, the resulting column submatrix is only rank 1, because the first and second columns are linearly dependent. In contrast, DCSS with (*k* = 2, *ϵ* > 0.2) will select the first and third columns, and the resulting DCSS column submatrix will be rank 2. Unlike the first three methods, DCSS takes into account the collinearity between columns in the selection procedure. If the DCSS error tolerance for this toy example is less than 0.2, DCSS will select all three columns.

We also mention two asides: first, in applications where the number of cells is far greater than the number of gene features (*n* > *d*), the method can instead be applied to **A**^*T*^ instead of **A** to filter cells instead gene features; second, the method can be modified to select columns for any rank-*k* subspace defined by *k* singular vectors of **A**, and not just the leading-k subspace (e.g. drop component 1 but include component 2). This could be useful when some of the leading singular vectors are highly correlated with batch or other confounding effects.

### 2.2 New bounds for DCSS

We derive a new spectral approximation bound (Bound 2.9) for the square of the submatrix **C** selected with DCSS and projected onto the rank-*k* subspace that best approximates **A**.

THEOREM 2.1 Let 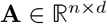 be a matrix of at least rank *k* and 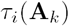 be defined as in Eqn. 1.1. Construct **C** following the DCSS algorithm described in Sec. 2.1. Then **C** satisfies,

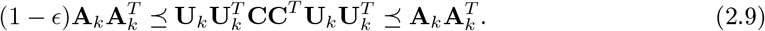

The symbol ⪯ denotes the Loewner partial ordering which is reviewed in Sec 6.1. Conceptually, the Loewner ordering is the generalization of the ordering of real numbers (e.g. 1 < 1.5) to Hermitian matrices. This bound means that after projection onto the rank-*k* subspace that best approximates **A**, **CC**^*T*^ is “close" to **AA**^*T*^ on that subspace. Statements of Loewner ordering are quite powerful; important consequences include inequalities for the eigenvalues and Euclidean distances. Some of the consequences of the Loewner ordering are reviewed in Sec 6.1. Bound 2.9 and the fact that **CC**^*T*^ ⪯ **AA**^*T*^ implies a bound on the Frobenius norm of *C*, a measure of the “size" of a matrix,

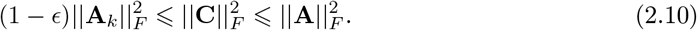

In the event that **A** is mean-centered, this means that the total empirical variance of **C** is bounded from below by (1 − *ϵ*) the variance in **A**_*k*_ and bounded from above by the total variance of **A**. The proof of Bound 2.9 and Bound 2.10 is included in Sec. 6.2.

One simple consequence of Bound 2.9 is the following bound,

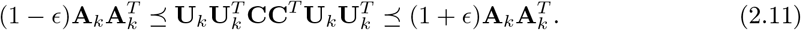

Bound 2.11 also holds for **C** selected by random sampling methods with *t* columns (see Sec. 6.3 for the theorem and proof). Thus, DCSS selects fewer columns with the same accuracy *ϵ* in Bound 2.11 for power-law decay in the rank-*k* subspace leverage scores when,

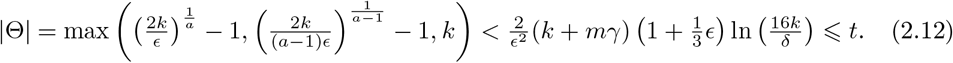

In this expression, *m* is the number of columns with zero rank-*k* subspace leverage score, *γ* is the minimum non-zero leverage score, and *δ* is the probability that Bound 2.11 fails to hold under random sampling.

## 3. RESULTS

We present two case studies where we compare DCSS to the simple thresholding methods of variance, count, and index of dispersion. We analyze the overlap in the selected columns. We also illustrate the effect of DCSS compared to the complete data for single-cell clustering.

### 3.1 Zeisel et al. (2015)

As a concrete illustration of the DCSS method, we focus on the genome-wide expression profiles of 3005 cells from the mouse somatosensory cortex and hippocampal CA1 region (Zeisel *et al.,* 2015) and the clustering workflow of Ntranos *et al.* (2016). The main contribution of Ntranos *et al.* (2016) is to perform clustering directly on the partition of reads into equivalence classes (ECs) rather than on a full quantification of reads into gene expression. ECs are a partition of reads into distinct classes, such that every read in a class maps to exactly the same set of transcripts (Nicolae *et al.,* 2011). This method is computationally scalable, comparable across scRNA-Seq experiments, and can be more accurate than clustering performed on a full quantification of reads into gene expression profiles (Ntranos *et al.,* 2016).

The Ntranos *et al.* (2016) data matrix **A** is 3,005 cells × 246, 981 EC counts. By the elbow method, we choose *k* = 5 for the DCSS workflow (Fig. 1a). We select an error tolerance of *ϵ* = 0.1. The rank-5 subspace leverage scores and the power-law fit for the top-scored 10, 000 ECs are shown in Fig. 1b. The column submatrix **C** has only 862 ECs, or approximately 0.3% of the total ECs. These ECs contain 42.3% of the reads. These 862 ECs map to 2,748 transcripts and to 1,642 genes. Table 1 contains the gene ontology term enrichment analysis (The Gene Ontology Consortium, 2015) on the genes corresponding to the DCSS (*k* = 5, *ϵ* = 0.1) ECs. Enrichments relevant for the brain include neuron part, neuron projection, and olfactory receptor activity. The enrichment analysis has an important caveat: because we map ECs to transcripts without positing an error model, there could be a high rate of false positives in the resulting transcripts and genes.

**Fig. 1:**
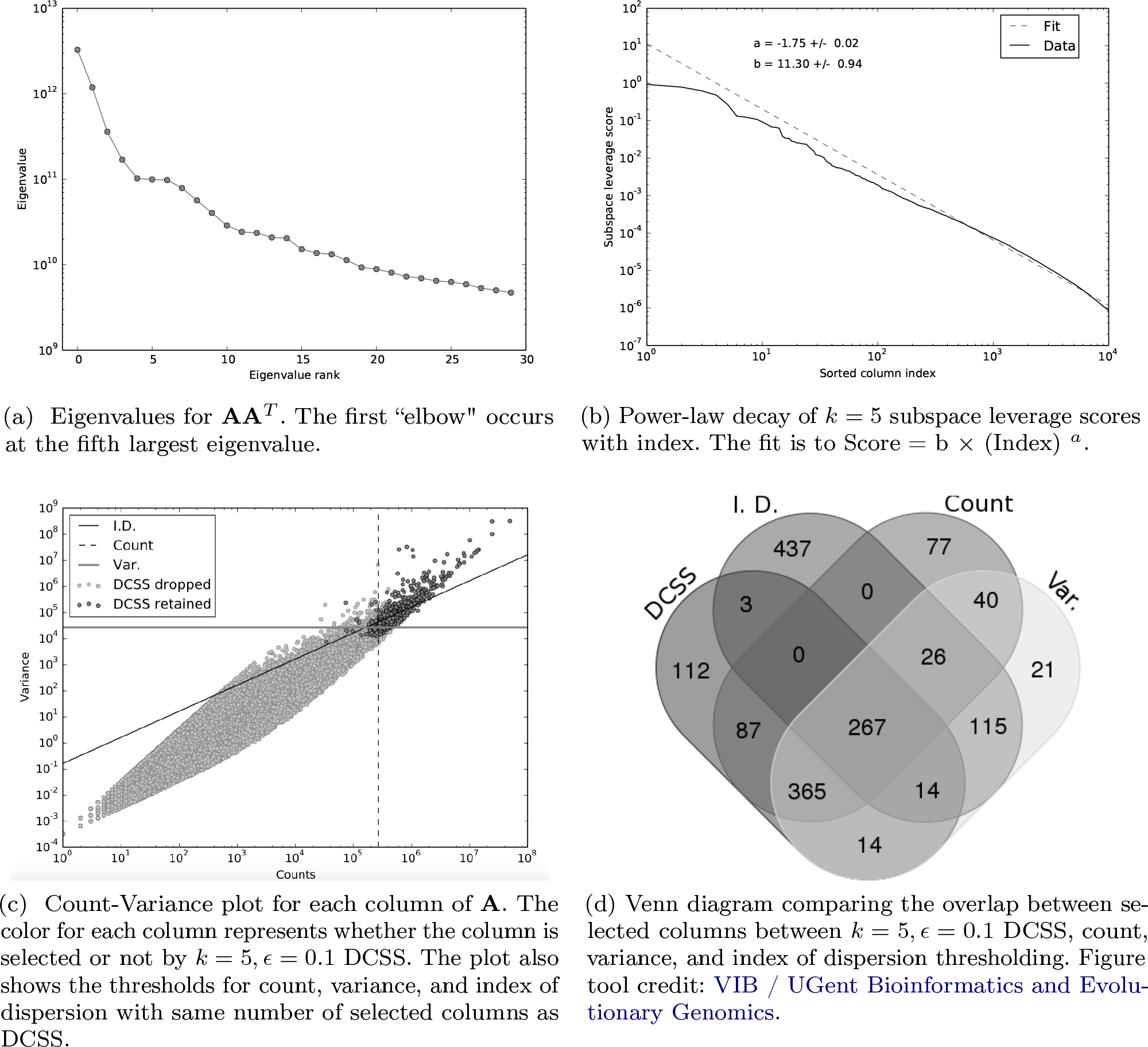
Figures for the Zeisel et *al.* (2015) and Ntranos et *al.* (2016) dataset.

**Table 1:** PANTHER overrepresentation test (release 20160715) with the GO Ontology database (release 2016-08-22) for the *k* = 5, *ϵ* = 0.1 DCSS 862 ECs mapped to 1,642 genes.

| Type | Gene ontology (GO) term | Bonferroni<br>p-value |
| --- | --- | --- |
| Biological process | cellular component organization (GO:0016043) | 1.12E-02 |
| Biological process | cellular component organization or biogenesis<br>(GO:0071840) | 8.01E-03 |
| Biological process | localization (GO:0051179) | 4.37E-02 |
| Cellular component | neuron projection (GO:0043005) | 4.52E-04 |
| Cellular component | neuron part (GO:0097458) | 8.24E-05 |
| Cellular component | cell projection (GO:0042995) | 8.36E-03 |
| Cellular component | cytoplasm (GO:0005737) | 1.59E-02 |
| Cellular component | intracellular part (GO:0044424) | 4.89E-02 |
| Molecular function | enzyme binding (GO:0019899) | 3.35E-02 |
| Molecular function | olfactory receptor activity (GO:0004984) | 1.30E-02 |

We compare DCSS to the three simple thresholding methods with the same number of columns in Fig. 1c and Fig. 1d. These figures show the similarities and differences in columns selected by the four thresholding methods. The simple thresholding methods have sharp boundaries in Fig. 1c, while the DCSS boundary is not linearly separable. The DCSS boundary approximately interpolates between the count and variance boundaries, and is most distinct from the index of dispersion boundary. Fig. 1d summarizes the overlap between selected columns in a Venn diagram. These figures illustrate that the DCSS method selects columns that are highly variable, have large counts, and frequently are over-dispersed; as such, the DCSS method is prescribed for quality control and to control technical variability.

The Ntranos *et al.* (2016) workflow for the Zeisel *et al.* (2015) dataset is to perform spectral clustering on pairwise Jensen-Shannon (JS) distances derived from the partition of reads into ECs. The spectral clustering clustering algorithm used is standard; the algorithm is to perform *k*-means clustering on the *k*-dimensional SVD projection of the normalized Laplacian of the symmetric similarity matrix **S**. The similarity matrix used for spectral clustering is *S*(**p**, **q**) = 1 − D_*JS*_(**p**, **q**), where D_*JS*_(**p**, **q**) is the JS distance between two probability mass functions p, q *ϵ* 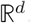. JS distances are well-suited to high-dimensional data, and provide more accurate clustering than *L_2_* distances on scRNA-Seq data (Ntranos *et al.,* 2016). For the Zeisel *et al.* (2015) data, the probability mass function for each cell is the vector of EC counts, normalized to sum to one. For the four thresholded workflows (DCSS, count, variance, and index of dispersion), the probability mass function for each cell is the subset vector of EC counts, normalized to one.

We evaluate the average spectral clustering classification error between the complete data and thresholded workflows, regarding the complete data workflow as the ground-truth. Since spectral clustering requires a random initialization for *k*-means, the average is over *T* =10 random initializations. Fig. 2 shows the average spectral clustering classification error rate for both two and nine spectral clusters for the workflow with the matrix **A** and the workflow with the column submatrix **C** for various *k*, *ϵ*. The different cells were curated into 47 subtypes by Zeisel *et al.*(2015), but we evaluate our method on courser-grained classifications because we have higher confidence in the spectral clustering ground-truth. Two spectral clusters identify neurons and non-neurons, while nine spectral clusters only loosely correspond to the nine major cell types. We also include the error for the three simple thresholding methods with the same number of columns as the DCSS method. We find that 0.3% of the total ECs give an error rate of 1.7% compared to the complete data for two clusters for *k* = 5, *ϵ* = 0.1 DCSS; only a small fraction of the gene expression profiles currently produced in scRNA-Seq experiments may be necessary to obtain the clusters reflecting cell types.

**Fig. 2:**
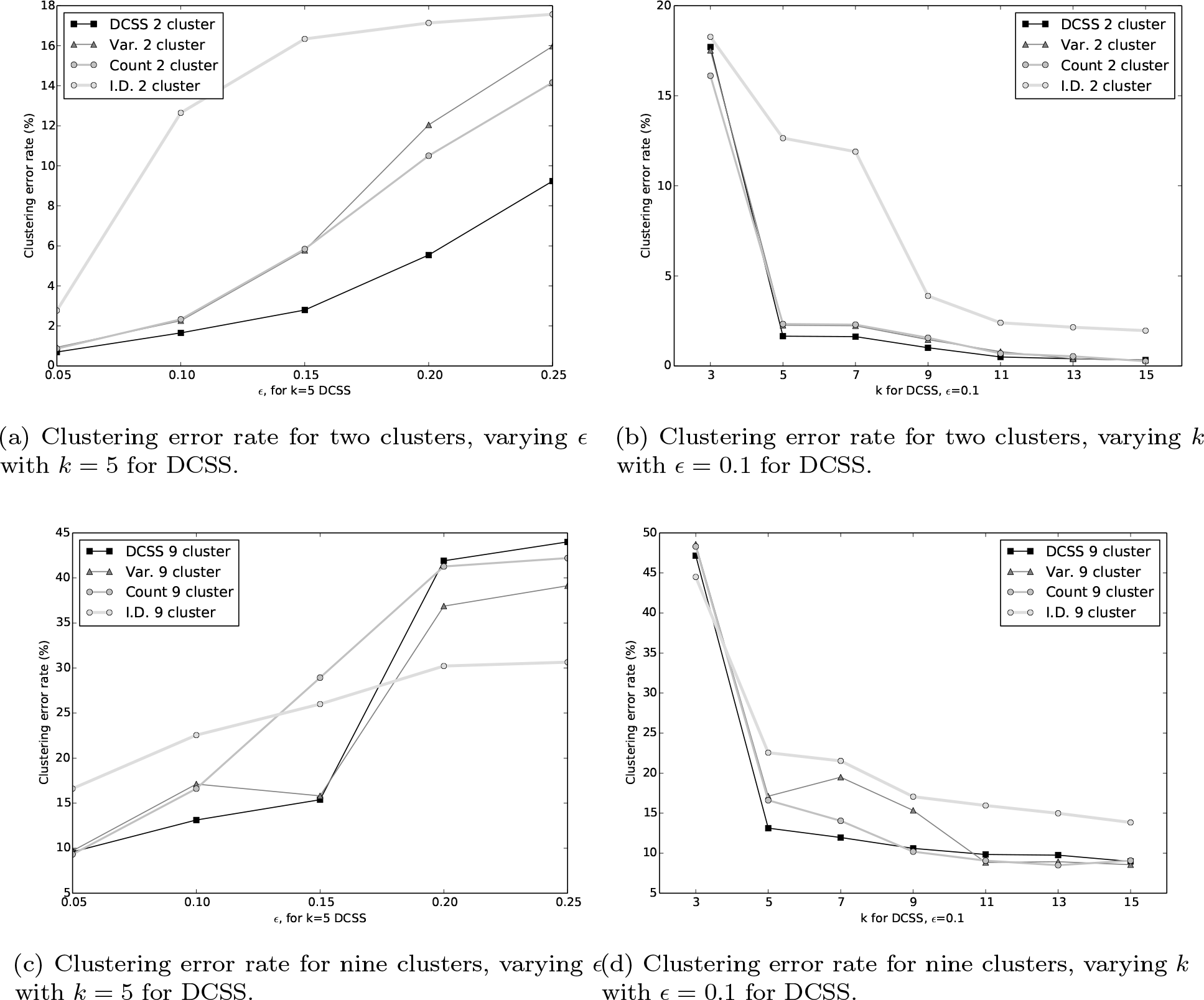
Average spectral clustering error for two and nine clusters for DCSS, count, variance, and index of dispersion threshoding for the Zeisel et *al.* (2015) and Ntranos et *al.* (2016) dataset.

### 3.2 *Paul* et al. *(2015)*

As a second application of the DCSS method, we focus on the genome-wide mRNA expression profiles of 4, 423 cells from mouse bone marrow myeloid progenitors (Paul *et al.,* 2015), and the *wishbone* trajectory workflow of Setty *et al.* (2016). The contribution of Setty *et al.* (2016) to scRNA-Seq is to use diffusion maps to identify components related to the development and maturation of cells, specifically myeloid and erythroid progenitors from hematopoietic stem and progenitor cells (HSPCs).

The Setty *et al.* (2016) data matrix for the (Paul *et al.,* 2015) dataset is **A** is 4, 423 cells × 14, 955 gene unique molecular identifier (UMI) counts. The Setty *et al.* (2016) workflow is quite involved. In brief, the *wishbone* algorithm creates a nearest-neighbor Euclidean distance graph. This graph is used to estimate all of the shortest path distances between a set of randomly sampled cells and the rest of the cells, and the shortest path distances are used to make the trajectory and branch assignments. The ***wishbone*** algorithm acts on a set of diffusion components which are selected for immune cell differentiation through a gene-set enrichment analysis. The diffusion components are calculated from the diffusion map of the similarity matrix derived from the Gaussian kernel of the 10-nearest-neighbor Euclidean distance matrix from the 15-dimensional PCA projection of the normalized UMI gene counts (Setty *et al.,* 2016).

We choose *k* = 14 for the DCSS workflow by the elbow method (Fig. 3a). We choose *k* = 14 rather than an elbow at a smaller *k* because the diffusion component workflow is sensitive to more components. We select an error tolerance of *ϵ* = 0.05. The rank-14 subspace leverage scores and the power-law fit for the top-scored 5, 000 genes are shown in Fig. 3b. The column submatrix **C** has 4, 693 genes, or approximately 31.4% of the total genes. These genes contain 90.4% of the UMI counts.

**Fig. 3:**
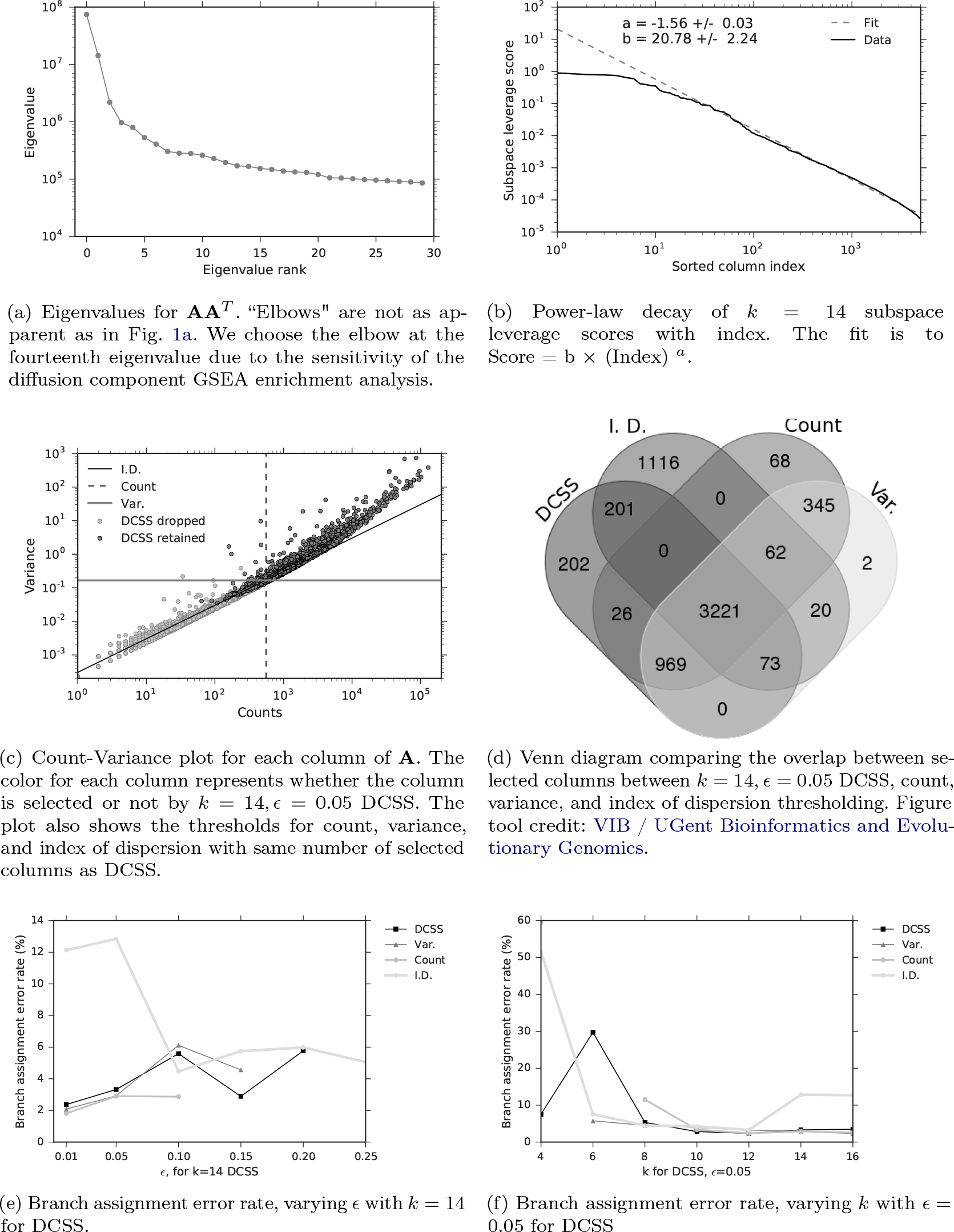
Figures for the Paul et *al.* (2015) and Setty et *al.* (2016) dataset.

We compare DCSS thresholding with *k* = 14, *ϵ* = 0.05 to the three simple thresholding methods with the same number of columns in Fig. 3c and Fig. 3d. The distribution of columns on the count-variance plots are qualitatively different between the Paul *et al.* (2015) data (Fig. 3c) and the (Zeisel *et al.,* 2015) data (Fig. 1c). This difference is expected due to the differences between ECs and gene UMI counts. Although the index of dispersion method is more differentiated from the other methods on the Paul *et al.* (2015) dataset, the behavior of the DCSS method in relation to the simple thresholding methods is similar between the datasets.

We calculate the average *wishbone* classification error between the two workflows, again regarding the complete data workflow as the ground-truth. Since the *wishbone* algorithm utilizes random sampling, the average is over *T* =10 *wishbone* branch assignments. The original *wishbone* analysis included only diffusion components 1 and 2. We additionally include diffusion component 4, since it is also enriched for immune cell differentiation according to the GSEA. For the Paul *et al.* (2015) dataset, *wishbone* assigns cells to three branches. Setty *et al.* (2016) used the behavior of four markers (CD34, Gata1, Gata2, and Mpo) to verify that the three branches correspond to HSPCs, myeloid progenitors, and erythroid progenitors, and the behavior does not change with the inclusion of component 4. Fig. 3 shows the average branch assignment classification error rate for the workflow with the matrix **A** and the workflow with the column submatrix **C** for various *k*, *ϵ*, and also the three simple thresholding methods with the same number of columns as the DCSS method for each *k*, *ϵ* point. Not all the thresholding methods successfully complete the *wishbone* workflow at large *ϵ*, due to the sensitivity of the diffusion component GSEA enrichment analysis, which we perform with keyword string matching. We find that for the *k* = 14, *ϵ* = 0.05 DCSS, 31.4% of the total genes give an error rate of 3.3% for three branch assignments compared to the complete data; this supports our conclusion that only a small fraction of the gene expression profile from scRNA-Seq experiments may be necessary to obtain meaningful cell-type classifications.

## 4. DISCUSSION

scRNA-Seq experiments allow researchers to probe the cell-specific heterogeniety in gene expression. Quality control and technical variability are significant challenges for scRNA-Seq experiments, and additionally the whole-genome expression profile is high-dimensional. In this note, we explore three existing simple thresholding schemes- count, variance, and index of dispersion- and propose a novel application of a thresholding scheme - DCSS- to select informative features and control quality and technical variability. We prove a bound on the “closeness" of the DCSS data submatrix to the complete data matrix (Eqn. 2.9), enlarging upon the existing set of error guarantees for DCSS (Papailiopoulos, Kyrillidis and Boutsidis, 2014), and illustrating the advantage of DCSS over the three simple thresholding schemes. Other advantages of DCSS include sensitivity to collinearity of features and covariance of cells. Since scRNA-Seq experiments are frequently used to cluster and classify cells, we choose the error rate for clustering and classification compared to the complete data as the evaluation metric for these thresholding schemes.

We present two case studies, the first on mouse cortex and hippocampus scRNA-Seq (Zeisel *et al.,* 2015; Ntranos *et al.,* 2016), and the second on mouse bone marrow scRNA-Seq (Paul *et al.,* 2015; Setty *et al.,* 2016). For the mouse cortex, the data matrix is cells × ECs, and only an incredibly small fraction of the ECs are necessary to obtain neuron and non-neuron cell clusters. For the mouse bone marrow, the data matrix is cells × genes, and only a small fraction of the genes are necessary to obtain HSPC, myeloid progenitor, and erythroid progenitor branch assignments. For both case studies, DCSS performs similarly to the simple thresholding schemes, in that it reduces the low abundance genes, maintains the most variable and over-dispersed genes. This supports our recommendation to use DCSS to control quality and technical variability. In both case studies, the error rate between the clustering computed with the complete expression profile and the reduced expression profile is small, suggesting that the clustering algorithms rely on a small subset of informative features.

## 5. SOFTWARE

The Python-package containing code to perform the methods described in the article can be found at https://github.com/srmcc/dcss_single_cell.git. The package also contains code to download the datasets used as examples in the article.

## ACKNOWLEDGMENTS

Research reported in this publication was supported by the National Human Genome Research Institute of the National Institutes of Health under Award Number [F32HG008713]. The content is solely the responsibility of the authors and does not necessarily represent the official views of the National Institutes of Health. SRM would like to acknowledge Ilan Shomorony and Robert Tunney for useful comments.

## 6. APPENDIX

### 6.1 Brief linear algebra review (Horn and Johnson, 2013)

*The singular value decomposition* (SVD) of any complex matrix **A** is **A** = **UΣV**^†^, where **U** and **V** are square unitary matrices (**U**^†^**U** = **UU**^†^ = **I**, **V**^†^**V** = **VV**^†^ = **I**), **Σ** is a rectangular diagonal matrix with real non-negative non-increasingly ordered entries. **U**^†^ is the complex conjugate and transpose of **U**, and **I** is the identity matrix. The diagonal elements of **Σ** are called the *singular values*, and they are the positive square roots of the eigenvalues of both **AA**^†^ and **A**^†^A, which have eigenvectors **U** and **V**, respectively. **U** and **V** are the *left* and *right singular vectors* of **A**.

Defining **U**_k_ as the first *k* columns of **U** and analogously for **V**, and **Σ**_k_ the square diagonal matrix with the first *k* entries of **γ**, then 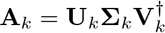is the rank-*k* SVD approximation to **A**, and **T** = **AV**_*k*_ = U_k_ X_k_ is a rank-*k* SVD truncation of **A**. Furthermore, we refer to matrix with only the last *n* − *k* columns of **U**, **V** and last *n* − *k* entries in **Σ** as **U**_\k_, **V**_\k_, and **Σ**_\k_.

The Moore-Penrose pseudo inverse of a rank *r* matrix **A** is given by 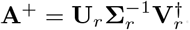.

The Frobenius norm ||**A**||_*F*_ of a matrix **A** is given by 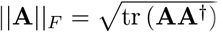. The spectral norm ||**A**||_2_ of a matrix **A** is given by the largest singular value of **A**.

The Eckart-Young-Mirsky theorem (Eckart and Young, 1936) states that, for **A** = **UΣV**^†^ the SVD of **A**, and **B** any complex matrix with compatible dimension to **A** and rank ≤ *k*,

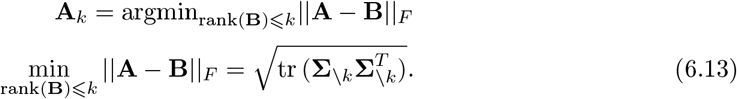

The minimizer **A**_k_ is unique if and only if σ_k+1_ ≠ σ_k_, where σ_*i*_ are the respective non-increasingly ordered singular values in Σ.

A square complex matrix *F* is *Hermitian* if **F** = **F**^†^ Symmetric positive semi-definite (S.P.S.D) matrices are Hermitian matrices. The set of *n × n* Hermitian matrices is a real linear space. As such, it is possible to define a *partial ordering* (also called a Loewner partial ordering, denoted by ⪯) on the real linear space. One matrix is “greater" than another if their difference lies in the closed convex cone of S.P.S.D. matrices. Let F, G be Hermitian and the same size, and *x* a complex vector of compatible dimension. Then,

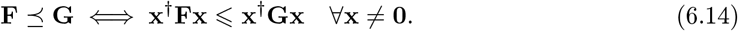

A few simple consequences of the Loewner partial ordering are as follows. If **F** is Hermitian and S.P.S.D., then 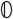⪯ **F**, where 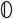 is the zero matrix.

If **F** is Hermitian with smallest and largest eigenvalues λ_min_(**F**), λ_max_(**F**), respectively, then,

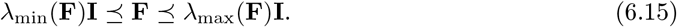

Let **F**, **G** be Hermitian and the same size, and let **H** be any complex rectangular matrix of compatible dimension. The *conjugation rule* is,

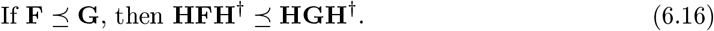

In addition, let λ_*i*_(**F**) and λ_*i*_(**G**) be the non-decreasingly ordered eigenvalues of **F**, **G**. Then

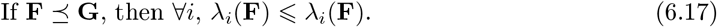

Since the trace of a matrix **F** is the sum of its eigenvalues, tr 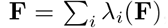, and the Loewner ordering implies the ordering of eigenvalues (Eqn. 6.17), the Loewner ordering also implies the ordering of their sum,

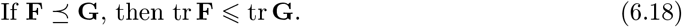

Let **F**_1_, **G**_1_, **F**_2_, **G**_2_ be Hermitian and the same size. Then if **F**_1_ ⪯ **G**_1_ and F_2_ ⪯ G_2_, then

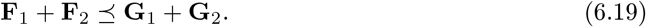

As a simple consequence of Eqn. 6.14, consider the real matrices **FF**^*T*^ and **GG**^*T*^, and the vector x which has a one in row *i* and a minus one in row *j*, and zeros elsewhere. The Euclidean distance between rows *i*, *j* with respect to **G** is *d_i,j_* (**G**):

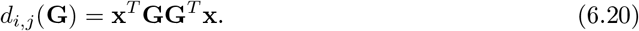

Thus, if **FF**^*T*^ ⪯ **GG**^*T*^, by Eqn. 6.14 with appropriate vectors, *d_i,j_*(**F**) ⪯ *d_i,j_* (**G**)∀*i*,*j*.

Furthermore, let F be Hermitian and dimension *n*, **U**_*k*_ be a semi-orthogonal rectangular matrix 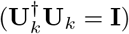 of compatible dimension *n* × *k*, 1 ≤ *k* ≤ *n*, and *λ*_*i*_(**M**) refer to the non-decreasingly ordered eigenvalues of a matrix **M**. Then the upper bound of the ***Poincaré separation theorem***states,

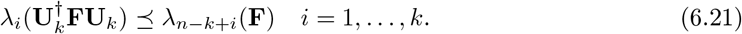

We will also use the von Neumann trace inequality. Let **F**, **G** be complex matrices of compatible dimension and minimum dimension *n*. Let σ_*i*_(**F**), σ_*i*_(**G**) be the respective non-increasingly ordered singular values. Then

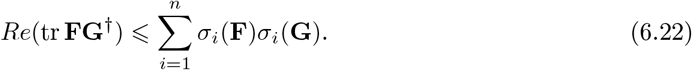

### 6.2 Proof of Bound 2.9

The upper bound (Bound 2.9) in Theorem 2.1 follows from the fact that 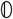 ⪯ **I** − **SS**^*T*^ and the conjugation rule (Eqn. 6.16),

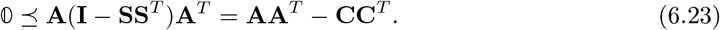

This upper bound is true for any column selection of **A**. A second application of the conjugation rule gives the upper bound in Bound 2.9.

For the lower bound (Bound 2.9), consider the quantity 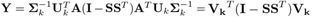. By the conjugation rule (Eqn. 6.16) on Eqn. 6.23, 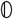 ⪯ **Y**, so **Y** is S.P.S.D. By the construction of DCSS (Eqn. 2.3) tr 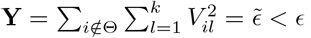, and because **Y** is S.P.S.D., λ_max_(**Y**) ≤ tr **Y**. By Eqn. 6.15 and the previous facts, **Y** ⪯ λ_max_(**Y**)**I** ⪯ *ϵ***I**. As a result of the conjugation rule applied to this upper bound,

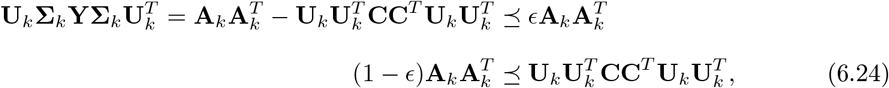

providing the lower bound of Bound 2.9.

For Bound 2.10, the lower bound of Bound 2.9 implies,

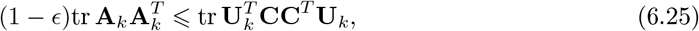

by Eqn. 6.18 and the cyclic property of the trace. Similarly, Eqn. 6.23 implies tr **CC**^*T*^ ≥ tr **AA**^*T*^. Since **U**_k_ is semi-orthogonal 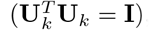, by Eqn. 6.21, every ordered eigenvalue of 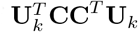 is smaller than its counterpart ordered eigenvalue of **CC**^*T*^. Since the trace is the sum of eigenvalues, this implies Bound 2.10,

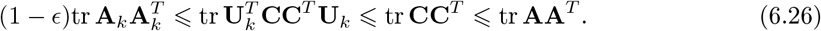

Note that if **A** is full rank and *k = rank*(**A**) = *n*, then Bound 2.9 becomes,

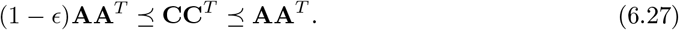

### 6.3 Proof of Bound 2.11 for random sampling

The following theorem pertains to a new spectral bound for the square **C** selected by rank-*k* subspace leverage scores and the random sampling procedure from Drineas, Mahoney and Muthukrishnan (2006).

THEOREM 6.1 Let 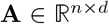 be a matrix of at least rank *k* and 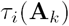 be defined as in Eqn. 1.1. Construct **C** by sampling *t* columns of **A**, reweighted to 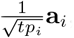, with probability 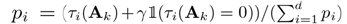, where 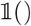 is the indicator function and *γ* is a small, positive, non-zero number 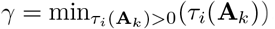. Let 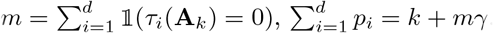. If the number of selected columns 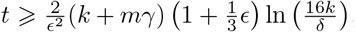 then with probability 1 − *δ*, the matrix **C** satisfies:

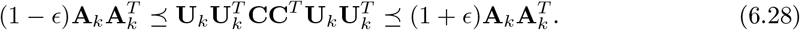

If **A** is full rank and *k* = *rank*(**A**) = *n*, then Bound 6.28 becomes,

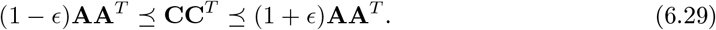

The proof of Theorem 6.1 is similar in structure to Theorem 3 in Cohen, Musco and Musco (2017). Theorem 3 in Cohen, Musco and Musco (2017) pertains to a different type of leverage score.

Consider the quantity 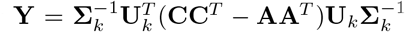. Note the sign change from Sec. 6.2. This can be rewritten as,

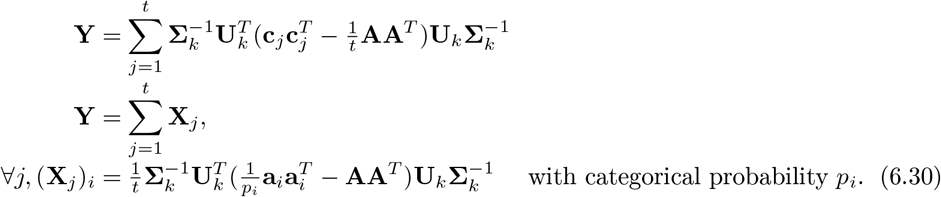

with categorical probability *p_i_*.

If ||**Y**||_2_ ≤ *ϵ*, then −*ϵ***I**⪯**Y**⪯ *ϵ***I**, and Bound 6.28 follows from this and the definition of **Y**. Thus, the proof of Bound 6.28 relies on showing that ||**Y**||_2_ ≤ *ϵ*. We use an intrinsic dimension matrix Bernstein inequality ((Tropp, 2015), Theorem 7.3.1), specialized to Hermitian matrices, to show that ||**Y**||_2_ is small with high probability. The Bernstein inequality requires that, for a finite sequence Y 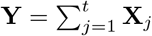 of random Hermitian matrices **X**_*j*_ of the same size,

1. 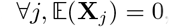,
2. ∀*j*,||**x**_*j*_||_2_ ≤ *L*,
3. and that 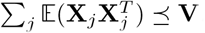.

Then, for 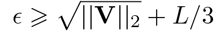,

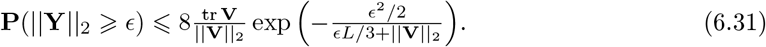

Requirement 1 is satisfied because,

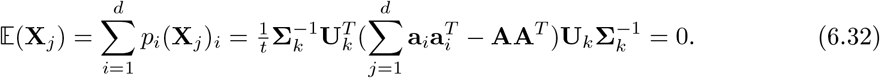

To show that requirement 2 is satisfied, we need the following fact:

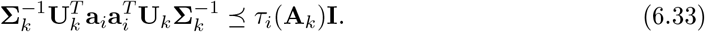

Eqn. 6.33 follows from the fact that for all y *ϵ* 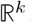,

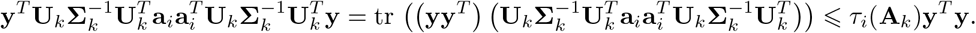

where the inequality comes from the Von Neumann trace inequality (Eqn. 6.22) applied to the product of two rank 1 matrices. Using Eqn. 6.33 in the definition of **X**_*i*_ gives,

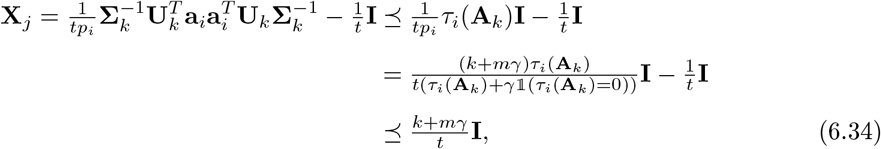

and 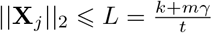 follows immediately.

To show that requirement 3 is satisfied, we compute directly,

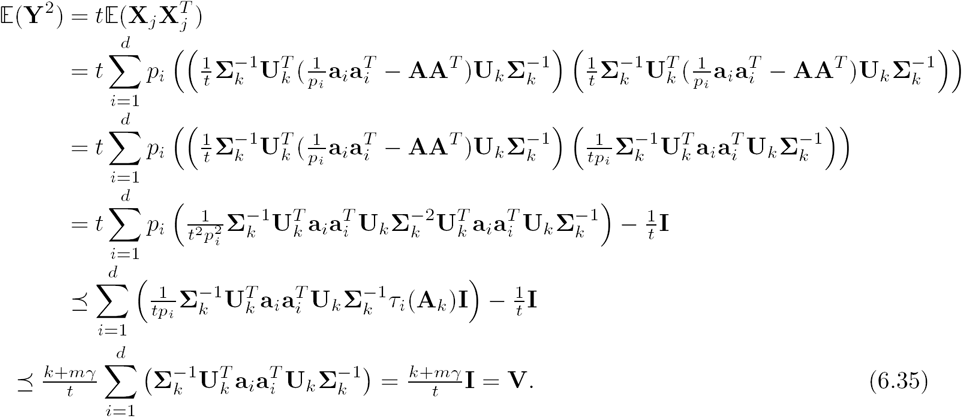

It follows immediately that 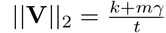 and tr 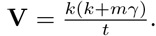

Then, for 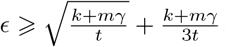,

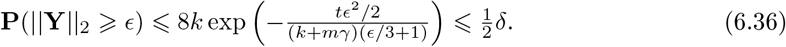

Solving for *t* as a function of *ϵ*, *δ*, and *γ* gives,

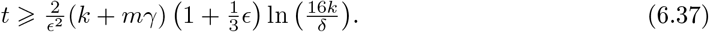

Bound 6.28 also holds for **C** selected by the DCSS algorithm, as a consequence of Bound 2.9. Thus DCSS selects fewer columns with the same accuracy for power-law decay for Bound 6.28 when |Θ| < *t*.

